# A Synthetic, Closed-Looped Gene Circuit for the Autonomous Regulation of RUNX2 Activity during Chondrogenesis

**DOI:** 10.1101/2021.04.20.440669

**Authors:** Gurcharan Kaur, Biming Wu, Sunjana Murali, Thomas Lanigan, Rhima M. Coleman

## Abstract

The transcription factor RUNX2 is a key regulator of chondrocyte phenotype during development, making it an ideal target for prevention of undesirable chondrocyte maturation in cartilage tissue engineering strategies. Here, we engineered an autoregulatory gene circuit (*cis*CXp-sh*Runx2*) that negatively controls RUNX2 activity in chondrogenic cells via RNA interference initiated by a tunable synthetic *Col10a1*-like promoter (*cis*CXp). The *cis*CXp-sh*Runx2* gene circuit is designed based on the observation that induced RUNX2 silencing after early chondrogenesis enhances the accumulation of cartilaginous matrix in ATDC5 cells. We show that the *cis*CXp-sh*Runx2* initiates RNAi of RUNX2 in maturing chondrocytes in response to the increasing intracellular RUNX2 activity without interfering with early chondrogenesis. The induced loss of RUNX2 activity in turn negatively regulates the gene circuit itself. Moreover, the efficacy of RUNX2 suppression from *cis*CXp-sh*Runx2* can be controlled by modifying the sensitivity of *cis*CXp promoter. Finally, we show the efficacy of inhibiting RUNX2 in preventing matrix loss in human MSC-derived cartilage under conditions that induce chondrocyte hypertrophic differentiation, including inflammation. Overall, our results demonstrated that the negative modulation of RUNX2 activity with our autoregulatory gene circuit enhanced matrix synthesis and resists ECM degradation by reprogrammed MSC-derived chondrocytes in response to the microenvironment of the degenerative joint.

## INTRODUCTION

Runt related transcription factor 2 (RUNX2) has a significant role in musculoskeletal development, including regulation of the chondrocyte phenotype during endochondral ossification^1,2^. Multiple intracellular pathways, including those mediated by SMADs^3^, β-catenin^4^, and p38^5^ signaling, are involved in hypertrophic maturation of chondrocytes during this process. These pathways are regulated by various extracellular signaling molecules, including bone morphogenetic proteins (BMPs)^6^, transforming growth factor beta (TGFβ)^7^, parathyroid hormone-related peptide (PTHrP)^8^, Indian hedgehog (IHH)^9^, fibroblast growth factor (FGF), and insulin like growth factor (IGF)^10^ as well as extracellular matrix (ECM) ligands such as hyaluronic acid^11^.

Synthetic regulation of the RUNX2 pathway has long been a target for cartilage tissue engineering, but its multifaceted functions in cell biology have led to mixed results. For example, blocking SMAD1/5/8 after chondrogenic differentiation of mesenchymal stem cells (MSCs) prevents terminal differentiation and mineralization while sustaining production of the articular cartilage structural macromolecules aggrecan and collagen type II^12^. Similarly, blocking the Wnt/β-catenin pathway during chondrogenic culture of MSCs suppresses hypertrophy and *in vivo* mineralization^13^. Combined strategies using dynamic loading or hypoxic culture of MSCs in hydrogels containing ECM ligands can also regulate hypertrophic pathways^14,15^. However, it has also been shown that complete inhibition or silencing of these and other RUNX2-related pathways will prevent or significantly delay chondrogenesis^12,16^. While these studies have investigated the role of various pathways that regulate RUNX2 gene expression and activity in MSCs, the direct effect of stage-specific suppression of RUNX2 activity on cartilage tissue engineering outcomes has yet to be explored.

Gene silencing through RNA interference (RNAi) allows straightforward loss-of-function studies in mammalian cells to help understand the molecular mechanisms that underpin cartilage development. A virally introduced inducible short hair-pin RNA (shRNA) system provides temporal and reversible control of protein expression so that its activity may be evaluated *in vitro* over long culture periods, such as those required for tissue engineering. These insights could help us develop targeted silencing protocols in cartilage tissue engineering applications, which has advantages over broad spectrum inhibition of intracellular pathways that drive or respond to RUNX2 activity. However, chondrogenesis and subsequent maturation of MSCs is not a homogenous process. Therefore, inducible RNAi systems cannot be used to address the stage-specific effects of RUNX2 inhibition in cell populations that are at various stages of maturation. Furthermore, RNAi systems cannot be used to study the direct effect of exogenous cues on RUNX2-driven pathways.

Advances in synthetic biology allow us to reprogram cells with new functions by site-specific editing of the genome or combining basic genetic modules with well-characterized functions in novel ways to create gene circuits. The reprogrammed cells are then capable of autonomously detecting and adapting to changes in their environment. The utility of these techniques has been demonstrated in the production of stem cells with autoregulatory resistance to inflammatory stimulus^17,18^. Autoregulatory gene circuits can be used to evaluate the impact of exogenous cues on RUNX2 activity levels as well as regulate the maturation of chondrogenic cells mediated by RUNX2.

In this study, we sought to engineer an autoregulatory, closed-loop gene circuit to negatively regulate RUNX2 activity in chondrogenic cells. Using an inducible RNAi system^19^, the effect of timing of RUNX2 interference on the retention of articular cartilage-specific structural macromolecules was first assessed. Based on the outcomes of these experiments, we then designed an RNA-based regulatory unit containing a partial *Col10a1*- like promoter region with tunable activity that drives expression of short hairpin RNA (shRNA) for RUNX2. Therefore, we constructed compact synthetic promoters *de novo* that contain multiple copies of RUNX2 *cis* enhancers upstream of the essential regulatory elements to initiate transcription. This design permits tuning of the gene circuit’s sensitivity to intracellular RUNX2 concentration as well as control of RUNX2 activity levels. We characterized our circuits with *in vitro* specificity and sensitivity assays using the ATDC5 cell line. We then tested the hypothesis that the level of RUNX2 activity can be tuned by varying the number of *cis*-enhancers incorporated into the promoter and that targeted disruption of RUNX2 activity could increase the production and retention of structural macromolecules in a cell-state-dependent manner. Finally, we demonstrate that the gene circuit resists the expression of hypertrophy markers and subsequent matrix degradation by human mesenchymal stem cell-derived chondrocytes (MdChs) under hypertrophic and inflammatory stimulus.

## METHODS

### Experimental Design

#### Part 1: Design and Assessment of the Inducible *shRunx2* System

The experiments to characterize the effect of cell state-specific RUNX2 suppression on chondrogenesis and accumulation of cartilage-specific structural macromolecules are outlined in Fig. 1. ATDC5 cells were first transduced with lentiviruses expressing a TetOn inducible *Runx2* shRNA. Stable polyclonal cell populations were then established via antibiotic selection. RUNX2 suppression was then induced using doxycycline at day 0, 4, 7, 14, and 21 during a four-week chondrogenic program in monolayer culture followed by gene and protein expression profiling.

**Figure 1.**
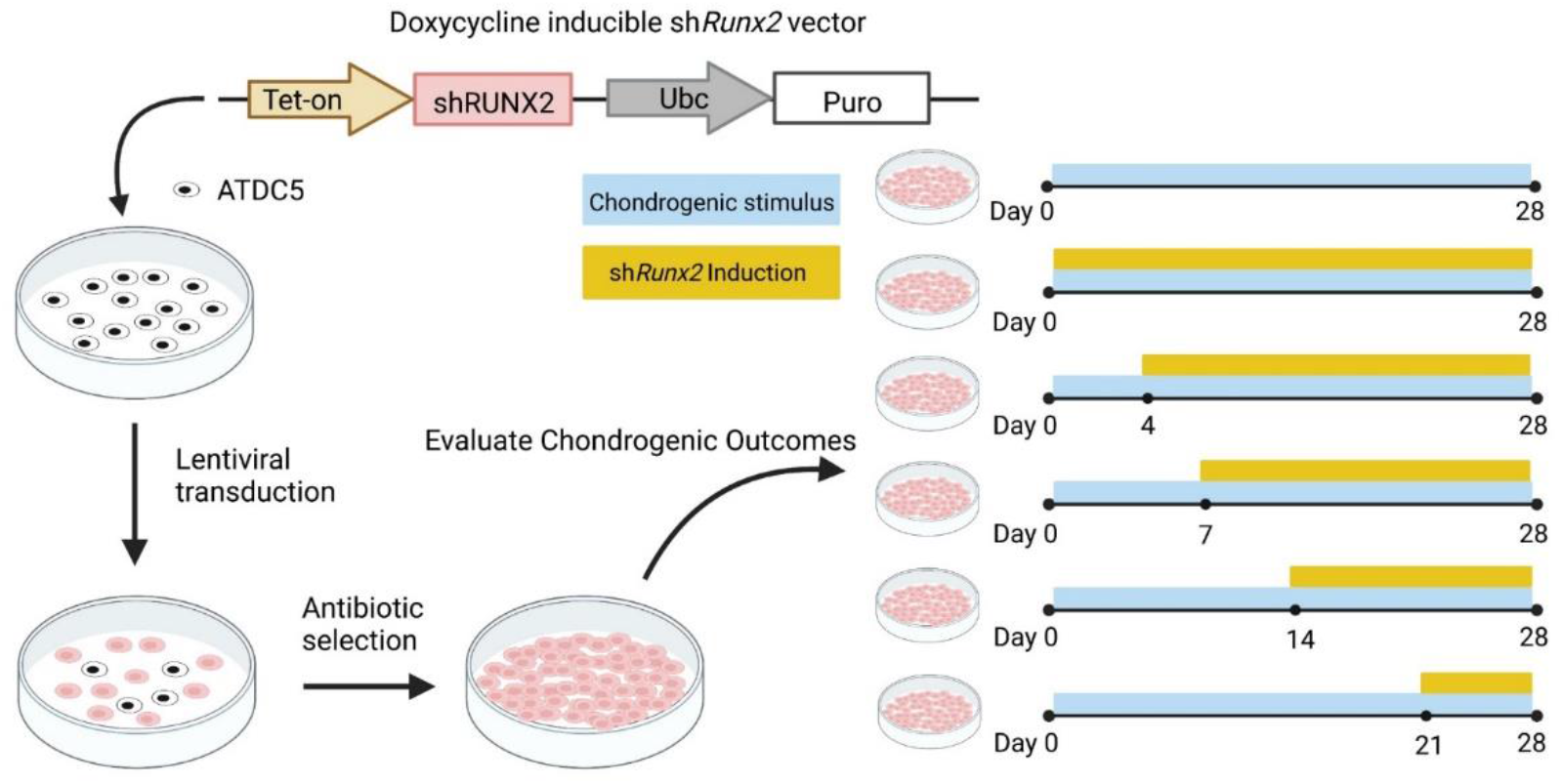
Experimental design of the inducible shRunx2 study.

#### Part 2: Design and Assessment of the Autoregulatory *shRunx2* Gene Circuit

To enable autoregulatory RUNX2 suppression and characterize its effect on chondrogenesis, we engineered a gene circuit that induces the expression of *Runx2* shRNA in response to increasing intracellular concentrations of active RUNX2 using a synthetic *Col10a1-like* promoter. Chondrogenesis of a polyclonal ATDC5 cell population stably expressing such a gene circuit was evaluated in both 3-week monolayer culture and 5-week pellet culture (Fig. 2).

**Figure 2.**
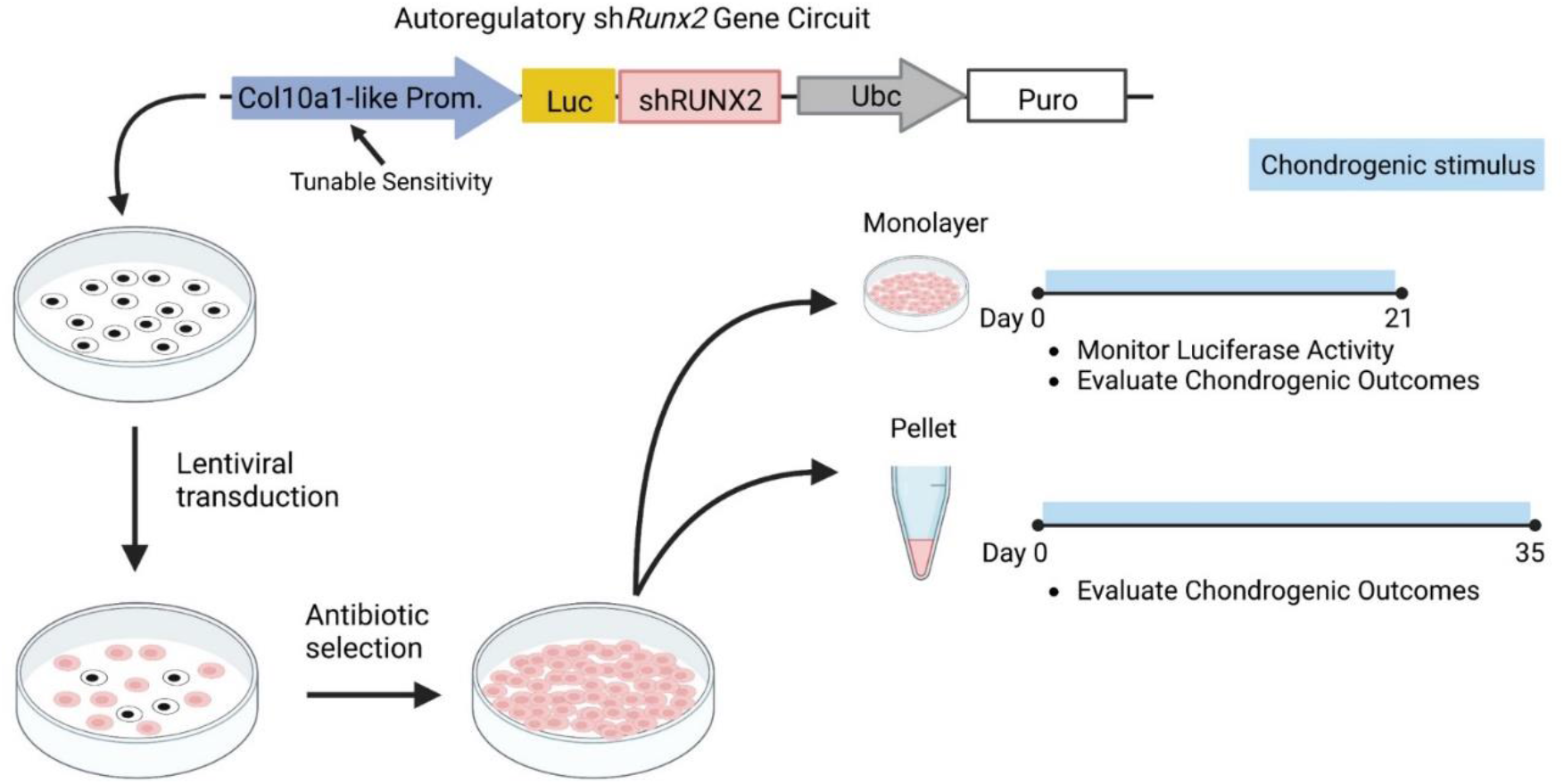
Experimental design of the autoregulatory *shRunx2* gene circuit study.

### Chondrogenic Cell Cultures

ATDC5 (Sigma; RRID:CVCL_3894) cells were used to model the transition through the different phases of endochondral ossification, as this chondroprogenitor cell line can recapitulate chondrogenic maturation in both monolayer (2D) and pellet (3D) cultures on an accelerated timescale^20–22^. After expansion, cells were transduced with the vectors described below. Chondrogenic behavior of these cells and their mineralization in response to exogenous phosphate treatment were carefully characterized in a previous study, which also describes the detailed chondrogenic culture protocol used in this study^23^. Briefly, chondrogenesis in 2D cultures of transduced or wild-type cells was initiated by addition of 1% ITS+ Premix (Corning) and 50 μg/ml L-ascorbic acid-2-phosphate upon 100% confluence. In 3D cultures, cell pellets containing 2.5×10^5^ cells were cultured in round-bottomed polypropylene 96-well-plates and media was changed every 2 days for 35 days.

Adult human bone marrow-derived MSCs (hMSCs) were plated at a density of 4,000 cells/cm^2^ and cultured in media containing (glucose DMEM + 10% FBS (Gibco qualified, lot# 1805387) + 1% Anti-anti+ 10ng/ml FGF-2 (Shenandoah, Cat# 100-146)). At ~80% confluency, cells were detached from plates with 0.25% trypsin/EDTA and subcultured. Experiments were performed using cells that underwent 12 population doublings (PD) as calculated from the formula PD = 3.32[log (final cell #) – log (0.2×10^6^)]. Pellets of transduced or wild-type cells were formed and cultured for either 21 or 28 days in chondrogenic media containing (DMEM, 4.5 g Glucose/L), 1% (v/v) Anti-Anti, 1% (v/v) ITS+ Premix (corning, cat# 354350), 1% Non-essential Amino Acids (Gibco, 11-140-050), 40 μg/ml L-proline (Sigma cat# P-0380), 50 μg/ml Ascorbic Acid 2-Phosphate, 0.1 μM Dexamethasone, 10 ng/ml TGFβ1 (Shenandoah, cat# 100-39)). For the inflammation studies, cell pellets were pre-cultured in chondrogenic media for 21 days. At day 21, the chondrogenic culture media was supplemented with 0.1ng/ml IL-1β (Shenandoah cat#100-167) in absence of TGFβ and dexamethasone to provide inflammatory stimulus. To induce hypertrophy, pellets were treated with 10nM triiodothyronine (T3) and 10mM beta glycerophosphate (βGP) starting on day 14 of chondrogenic culture^24^.

### Lentiviral Vector Design and Synthesis

#### Doxycycline Inducible shRunx2 Vector

The inducible system that drives RUNX2 knockdown was modified from the pINDUCER lentiviral toolkit originally developed by Meerbrey *et al*.^19^. pINDUCER13 was acquired from Addgene (plasmid # 46936) for further modification as such a lentiviral vector conveniently contains a Tet-on inducible cassette that co-expresses luciferase and shRNA sequence as well as a cDNA sequence of a constitutively active puromycin resistance gene (Fig 3a). Three shRNA sequences targeting murine/human *RUNX2* from RNAi Codex^25^ were screened for RUNX2 knockdown via western blotting and *shRunx2* sequence #294717 (5’-TGCTGTTGACAGT-GAGCGCCGAATGGCAGCACGCTATTAATAGTGAAGCCACAGATGTATTAATAGCGT-GCTGCCATTCGATGCCTACTGCCTCGGA-3’) was further selected for downstream vector synthesis.

**Figure 3.**
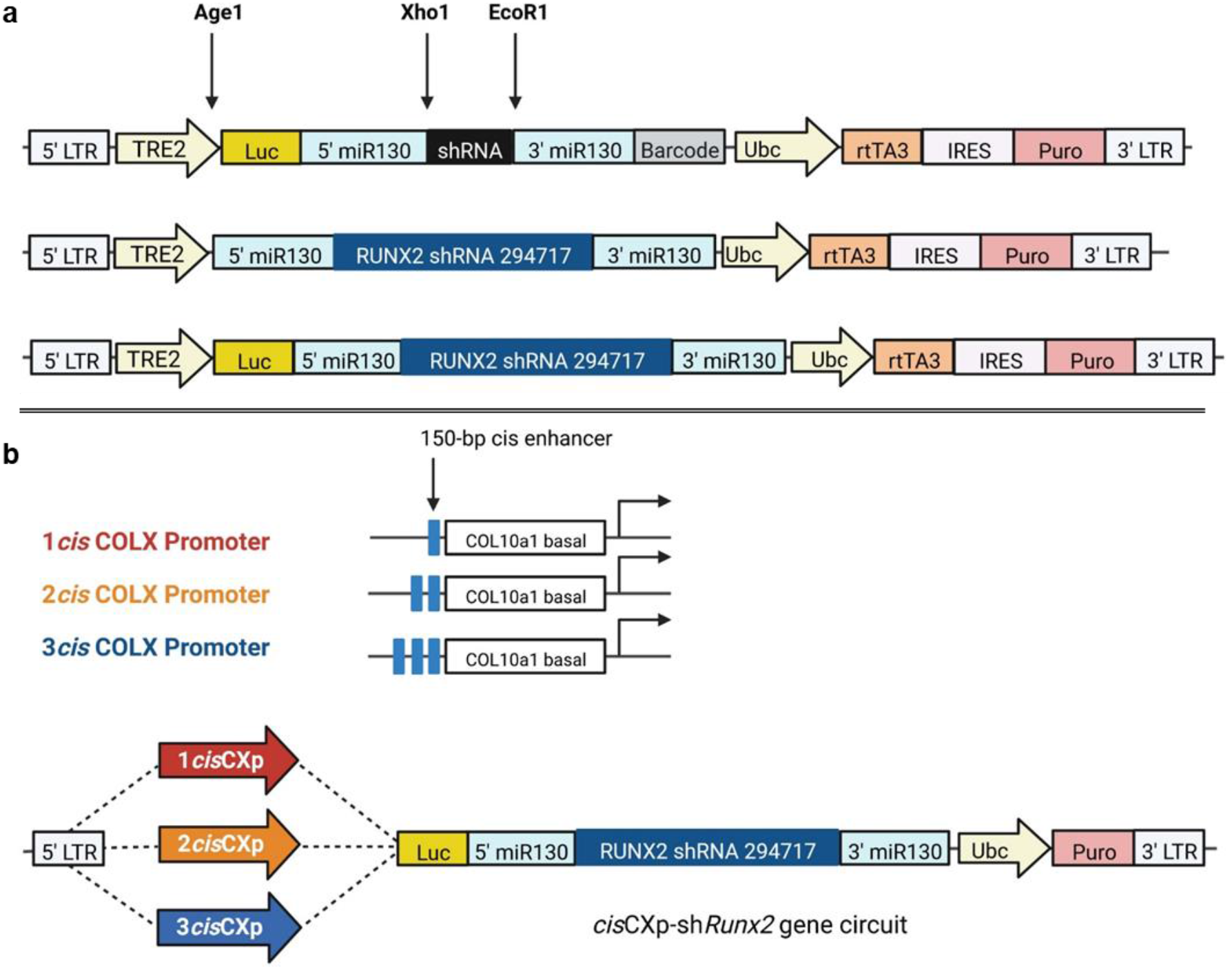
Diagrams of pINDUCER13, TetOn inducible *shRunx2* vectors and CXp promoter cisCXp-sh*Runx2* gene circuits. **(a)** TRE2, tetracycline response element 2 promoter; Luc, luciferase; Ubc, Ubiquitin C promoter; rtTA3, reverse tetracycline-controlled transactivator 3; Puro, puromycin resistance. Diagram of pINDUCER13 is adapted from Meerbrey *et al*^19^. (b) 1-, 2-, and 3cisCXp promoters contains one, two, and three copies of 150bp *Col10a1 cis*-enhancer upstream of *Col10a1* basal promoter.

Two inducible *shRunx2* vectors (TetOn-Luc-sh*Runx2* & TetOn-sh*Runx2*) were created. TetOn-Luc-sh*Runx2* was first created by sub-cloning PCR-amplified *shRunx2* #294717 into TetOn-inducible cassette of pINDUCER13 via XhoI/EcoRI sites. To remove the luciferase cDNA from TetOn-Luc-sh*Runx2*, miR30-shRunx2-miR30 sequence was PCR amplified with the addition of an AgeI site upstream of the 5’ end. The PCR product was subsequently sub-cloned into the TetOn-Luc-sh*Runx2* vector to replace the existing Luc-miR30-shRunx2-miR30. In addition, two scramble vectors (TetOn-Luc-scramble & TetOn-scramble) were synthesized to serve as experimental controls (Table S1). Expression of short hairpin sequences was induced by the addition of (0.5μg/ml) of doxycycline (Dox) beginning on the day indicated and continued to the end of culture.

#### Autoregulatory cisCXp-shRunx2 gene circuit

The autoregulatory *cis*CXp-sh*Runx2* gene circuits rely on a *Col10a1-like* promoter to drive RUNX2 knockdown via expression of *shRunx2* sequence. *Col10a1-like* promoters with a different number of *cis* enhancers were first reported by Zheng *et al*.^26^; single copies of 1*cis*CXp promoter which consists of one *cis* enhancer (−4296 to −4147 bp) upstream of the *Col10a1* basal promoter (−220 to 110 bp) were acquired from IDT technology (Fig. 3b). To create 1*cis*CXp-*shRunx2*, the 1*cis*CXp promoter was PCR amplified using corresponding primers (Table S2) so that both 5’- and 3’-ends of the reaction products have an overlap of >30 nucleotides with the backbone of TetOn-Luc-sh*Runx2* vector after TRE2 promoter was removed by restriction digestion with NheI/AgeI. The linearized Luc-shRunx2 vector and 1*cis*CXp promoter were further assembled together using Gibson Assembly Kit (NEB). A conceptual schematic of the gene circuits function is shown in Fig. 4.

**Figure 4.**
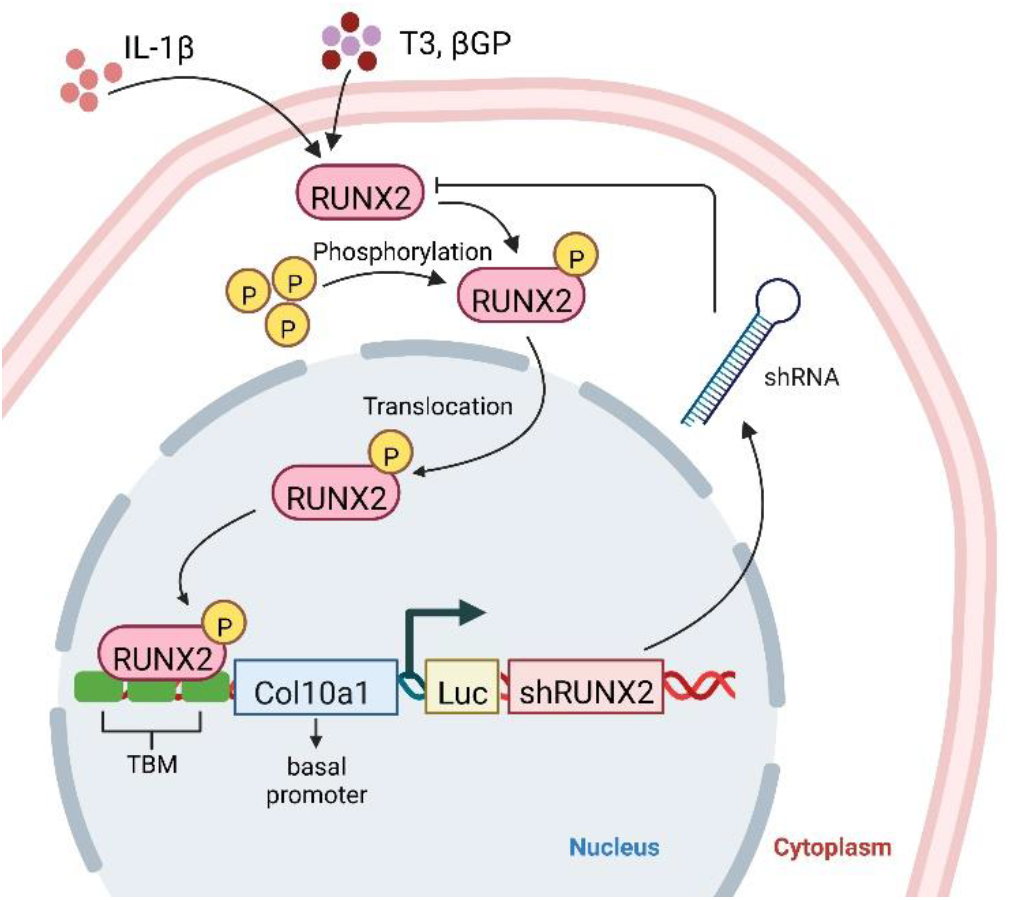
Schematic of the intracellular synthetic negative-feedback regulation of RUNX2. Maturation signals like T3, βGP and IL-1β upregulate RUNX2 expression which binds to the transcription factor binding motifs (TBM) in the gene circuit to induce the expression of shRUNX2, thereby establishing a negative-feedback loop that suppresses RUNX2 activity, allowing MdChs to resist maturation during neo-cartilage formation.

To create 2*cis*CXp-*shRunx2*, single copies of 150-bp *cis* enhancer were acquired from IDT technology. PCR amplifications of both the *cis*-enhancer and 1*cis*CXp promoter were performed using corresponding primers (Table S3) so that overlaps of >30 nucleotides are created between the reaction products to support the sequential assembly of linearized Luc-shRunx2 vector, *cis* enhancer, and 1*cis*CXp. Similarly, 3*cis*CXp-*shRunx2* was assembled from the three components described above as well as an additional PCR product that inserts an extra copy of *cis* enhancer in between the first *cis* enhancer and 1*cis*CXp promoter. Correct cloning was confirmed by Sanger sequencing at University of Michigan Sequencing core.

### Lentiviral Transduction and Stable Selection

To establish ATDC5 cell lines that express the vectors of interest, lentiviral supernatant of chosen plasmid vectors was produced by the University of Michigan Vector Core. Twenty-four hours prior to transduction, cells were plated in individual wells of a 6-well plate at a density of 10000 cell/cm^2^. Proliferating cells were transduced at multiplicity of infection (MOI) = 1 with lentiviral supernatant of TetOn-sh*Runx2*, *cisCXp*-Luc-sh*Runx2* gene circuits, and corresponding scramble vectors in the absence of serum for 48 hours. Polyclonal populations of cells stably expressing the chosen vectors were subsequently selected with continuous treatment of puromycin (2μg/ml) for 10-14 days for ATDC5 cells.

For human MSCs, 10,000 cells/cm^2^ were plated in individual wells of 6-well plate and transduced at MOI = 5 with lentiviral supernatant of *cisCXp*-Luc-*shRUNX2* gene circuits for 24 hours. Cells stably expressing the vectors were selected by treatment with 1μg/ml puromycin for 4 days.

For both cell types, cells were subcultured for one additional passage prior to each experiment as described above.

### Luciferase Assay

Luciferase activity of 2D and 3D cultures in multi-well plates was measured once daily beginning on day −1 or 0. Thirty minutes prior to each measurement, concentrated D-luciferin stock (1mg/ml) was added to existing culture media to achieve a final concentration of 150 μg/mL and gently mixed. Cultures containing luciferin were then incubated at 37°C for 30 minutes before measured using SYNERGY H1 microplate reader (BioTek). Each sample was measured three times (1 second/read) and the means were normalized to the day −1 values and expressed as relative luminescence units (RLU). Media was replaced in measured cultures immediately after reading.

### Biochemical Analysis

To assess the accumulation of cartilage-specific matrix in both 2D and 3D cultures, the sGAG levels were quantified using thebDMMB assay, as previously described^23,27^. Specifically, differentiated cell masses from 2D cultures or pellets from 3D cultures were washed with ice-cold PBS before being digested with 1mg/ml proteinase K in 200-500μl ammonium acetate (0.1M) at 50°C for 16 hours. sGAG content in the digested lysate of each sample was subsequently determined by comparing its DMMB reading to a standard curve. The DNA content of each sample was measured using Hoechst 33258 dye (Sigma) to normalize its corresponding sGAG content^28^.

### Gene Expression Analysis

Analysis of gene expression was performed as previously described^4^. Total RNA of both 2D and 3D cultures were extracted using TRI Reagent^®^ RT (Molecular Research Center). Cell masses from each individual well of 12-well plate were collected as one 2D sample. Four pellets were combined to generate 1 3D sample. Extracted RNA were reverse-transcribed into cDNA using High-Capacity cDNA Reverse Transcription Kit (Life Technologies). Relative expression levels were calculated as *x* = 2^-ΔΔ*C*^_*T*_, in which ΔΔ*C_T_* = Δ*C* − Δ*E*, 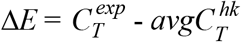 at the time-point of interest, 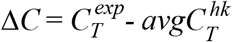 at day 0, and 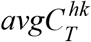 is the average *C_T_* value of two housekeeping genes (*Hprt* and *Ppia*). The forward and reverse primer sequences for all genes are listed in Table S2.

### Histological Analysis

Monolayer cultures were washed twice with PBS and fixed in 70% ethanol at room temperature for one hour. Fixed pellets were washed with 70% ethanol, embedded in paraffin wax, and then 7 μm sections were taken. Sections were deparaffinized in xylene, rehydrated in increasing dilutions of ethanol, stained with Alcian blue (1% in 3% Acetic Acid, Poly Scientific) for 30 minutes, and then counter-stained with Fast Nuclear Red. Stained sections were imaged using bright field microscopy.

### Immunohistochemistry

The pellet sections were deparaffinized in xylene and rehydrated in successive dilutions of ethanol. Slides were then blocked in 3% H_2_O_2_ for 10 minutes. This was followed by Antigen retrieval with heated Retrievagen A (BD Biosciences, Cat# 550524) for 20 minutes. Once cooled to room temperature, the samples were then permeabilized using 0.5% TBS-TX100, blocked with 10% Goat serum, 1% BSA in 0.1% TBST for 1 hour at room temperature, and then incubated overnight in either RUNX2 (ABclonal Cat# A2851; 1:100 dilution), COLX (ABclonal Cat# A6889; 1:100 dilution), MMP13 (ABclonal Cat# A16920; 1:200 dilution) or ACAN (ABclonal Cat# A8536; 1:500 dilution) primary antibody solution in 10% Goat serum in 0.1% TBS-TX100 at 4°C. After washing, samples were incubated with Goat Anti-Rabbit IgG (H+L) Alexa Fluor 488 secondary antibody (Invitrogen, Cat#A11034; 1:500 dilution in 10% Goat serum in 0.1% TBS-TX100) followed by 300nM DAPI solution. Lastly, the slides were cover-slipped with anti-fade mounting media.

### Western Blot Analysis

Whole cell extracts of 2D cultures from one well of a 6-well plate were prepared using RIPA Lysis and Extraction Buffer (Thermo Scientific) supplemented with protease inhibitor cocktail (Sigma) using the Micro Tube Homogenizer (Thermo Scientific). Homogenized lysate was rotated at 4 °C for one hour and centrifuged at 12,000*g* for 10 minutes to remove cellular debris. Total protein content within each sample was determined with the Pierce BCA Protein Assay Kit (Pierce). Proteins (5-15 *μ*g) were separated on a 10% NuPAGE Bis-Tris Protein Gel and then transferred to a polyvinylidene difluoride membrane (Millipore). Membranes were blocked with 5% BSA made up in Tris-buffered saline-Tween 20 (TBS-T, 0.1% Tween 20) for 1 hour at room temperature. Following blocking, membranes were incubated in TBS-T, 0.1% Tween 20 and 5% BSA overnight at 4 °C with anti-Runx2 antibody (RUNX2 D1H7 Rabbit mAb, Cell Signaling Technology, 84866 1:2000) and anti-β-actin antibody (Rabbit Anti-beta Actin, Abcam, ab119716, 1:5000). Secondary incubation was performed at room temperature for 1 hour using WesternSure^®^ Goat anti-Rabbit HRP (LiCor, 926-80011 1:20000). Positive staining was visualized using the LiCor WesternSure^®^ PREMIUM Chemiluminescent Substrate and quantified using the LiCor Image Studio.

### Statistics

Statistical analyses were performed in GraphPad Prism version 7 for Windows (GraphPad Software, La Jolla California USA) using one or two-way analysis of variance followed by Tukey’s post-hoc test. P-values less than 0.05 were considered statistically significant. All errors bars are the mean ± the standard deviation.

## RESULTS

### Part 1: Assessment of the Inducible *shRunx2* System

#### Constitutively Silencing of RUNX2 Inhibits Chondrogenesis

To examine the effect of constitutive RUNX2 silencing on *in vitro* differentiation of chondroprogenitors, we induced chondrogenesis in monolayer cultures of ATDC5 cells stably expressing TetOn-sh*Runx2 (shRunx2 cells)* in presence of doxycycline (Dox) and evaluated matrix accumulation and expression levels of chondrogenic markers after 14 days. In cultures treated with Dox, RUNX2 protein was depleted in sh*Runx2* cells by day 7 and remained downregulated through day 14 (Fig. 5a). With Dox treatment, there was also no detectable amounts of matrix accumulation in sh*Runx2* cells by day 14, while the scramble controls had successfully formed Alcian blue-positive nodules (Figs. 5b and c). The expression of mRNA for *Acan* and *Col2a1* was also significantly inhibited with Dox treatment of *shRunx2* cells (Fig. 5d), suggesting that constitutive RUNX2 suppression interfered with early chondrogenesis.

**Figure 5.**
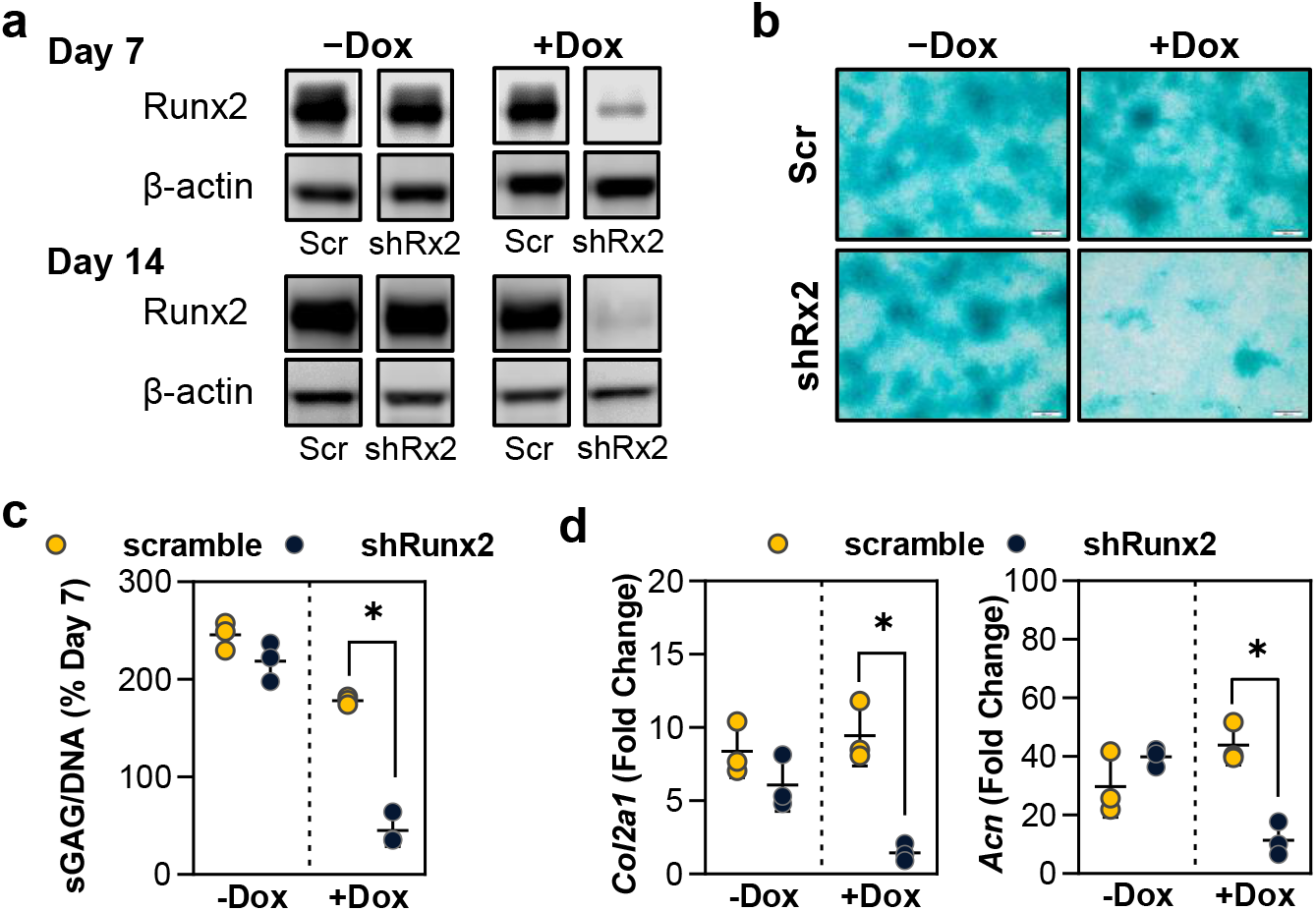
Effects of constitutively active RUNX2 silencing on early chondrogenesis. ATDC5 cells expressing Tet-on sh*Runx2*/scramble were differentiated in the presence (+) or absence (-) of 0.5 μg/ml doxycycline for 14 days. (a) Western blot analysis of RUNX2 and β-actin at day 7 and 14. Alcian blue staining (b) and % sGAG accumulation relative to samples at 7 days without Dox (c) at day 14. (d) Fold change in mRNA expression for aggrecan and collagen type II at day 14. Polyclonal populations were established by combining selected cells from two independent transduction experiments (two viral batches, n=3). All data represented as mean ± SD. Significant difference is indicated by * p<0.05.

#### Stage-dependent Effect of RUNX2 Silencing on Chondrogenesis

RUNX2 suppression in chondroprogenitors as they mature through the stages of endochondral ossification was examined by adding Dox continuously starting on the following time-points of culture that were found to correlate with specific stages of chondrocyte maturation, as determined in preliminary studies (Fig. 6c and Fig. S1): minimum RUNX2 protein expression (D4), just prior to rapid increase in RUNX2 protein expression (D7), peak gene expression of *Col10a1* (D14)^25^, and the peak of RUNX2 protein expression (D21). These were compared to cultures that received no Dox or were treated with Dox from day 0 (D0). Dox induction in monolayer suppressed RUNX2 protein levels by 20-60% in all cultures by day 28 (Fig. 5b). When RUNX2 suppression was induced from Day 4, 7, and 14, matrix accumulation was increased by 1.9-fold, 2.14-fold and 1.75-fold respectively on day 28 relative to scramble controls and the no Dox group (Fig. 6c). There was no difference between the scramble and no Dox groups.

**Figure 6.**
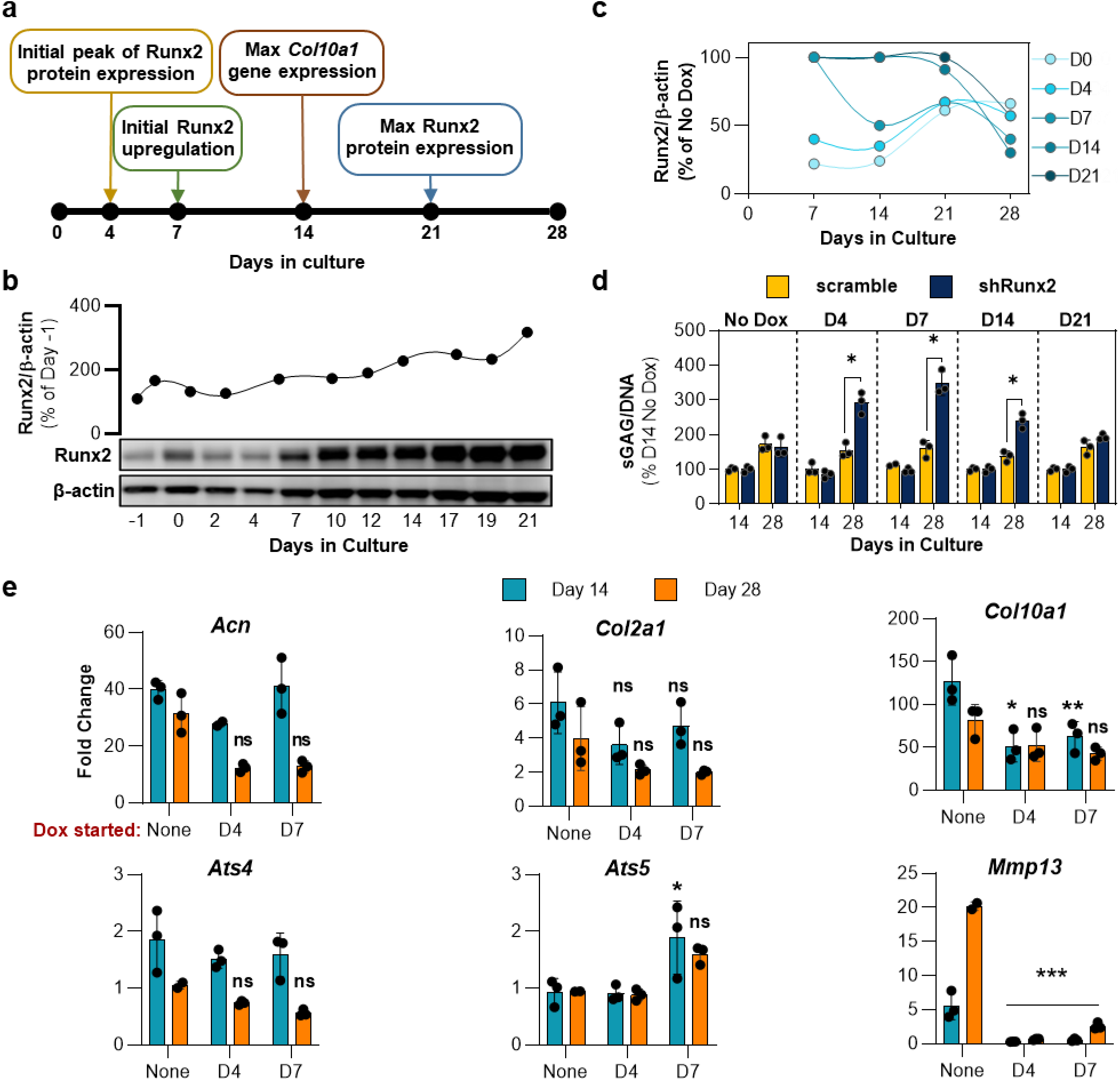
Effects of RUNX2 silencing at different maturation stages of chondrogenesis. (a) The time-course of cell-state transitions and relative Runx2 protein expression in monolayer cultures of ATDC5 cells. (b) Runx2 protein band intensity normalized by β-actin. The plot shows the percent increase in protein expression compared to just prior to induction of chondrogenesis (day −1). (c) Relative RUNX2 protein expression in TetOn-sh*Runx2* cells in response to Dox treatment from different time points of chondrogenic differentiation relative to the No Dox controls. (d) Fold change relative to D14 No Dox group in sGAG accumulation normalized to DNA content, after 14 and 28 days in culture. Data represented as mean ± SD. Polyclonal populations were established by combining selected cells from two independent transduction experiments (two viral batches, n=3). (e) Effects of delayed RUNX2 silencing on gene expression of chondrogenic markers and matrix degrading enzymes. Quantification of mRNA expression of Acn, Col2a1, Col10a1, Adamts4, Adamts5, and Mmp13 at day 14 and 28 in cells with inducible expression of shRUNX2 beginning on D4 or D7. Polyclonal populations were established by combining selected cells from two independent transduction experiments (two viral batches, n = 3) Significant difference between TetOn sh*Runx2* and scramble groups at each time point is indicated by * p<0.05.

We next examined the effect of Runx2 suppression beginning on D4 or D7 on genes associated with chondrocyte hypertrophy, ECM production, and ECM turnover (Fig 5e). Runx2 suppression reduced expression of the hypertrophy marker *Col10a1* by 2-fold at 14 days of culture. This level was maintained through day 28. Expression of *Mmp13*, another significant marker of hypertrophy, was downregulated 5-20-fold by Runx2 suppression independent of when Dox treatment began. When RUNX2 was suppressed beginning on day 4 or day 7, gene expression of the chondrogenic ECM markers aggrecan and collagen II were not significantly reduced compared to controls that did not receive Dox. Runx2 suppression beginning at day 4 had no effect on *Adamts5* gene expression while delaying suppression until day 7 lead to increased expression of this metalloproteinase. Suppression beginning at either time point had no effect on *Adamts4* expression.

This inducible study indicates that suppression of RUNX2 at specific points of chondrocyte maturation has a differential impact on matrix accrual of cartilage-specific structural macromolecules and regulation of metalloproteinases expression. Therefore, persistent and broad-spectrum silencing of RUNX2 could have deleterious effects on chondrogenesis and matrix production in heterogeneous populations of chondroprogenitors at various stages of maturation. This makes it difficult to use inducible silencing to optimally target its suppression. Next, we engineered an autoregulatory genetic modification that would allow the cells to independently regulate their response to exogenous cues that raise intracellular RUNX2 levels.

### Part 2: Design and Assessment of the Autoregulatory *shRunx2* Gene Circuit

#### Engineering of a Synthetic Col10a1-like Promoter

To engineer a RUNX2-responsive promoter that is specific to chondrogenic cells, we assembled the 150bp RUNX2-binding *cis*-enhancer of the *Col10a1* promoter upstream of a truncated basal sequence from the same promoter (n*cis*CXp; Fig. 3). To demonstrate that this promoter is only active in chondrogenic cells undergoing hypertrophic maturation, a promoter containing 1 *cis*-enhancer (1*cis*CXp) was used to drive the expression of luciferase or enhanced green fluorescent protein (eGFP; Fig. 7a). Luciferase activity mirrored the time-course of *Col10a1* gene expression (Fig. 7b) in chondrogenic ATDC5 cells: minimal until day 6 followed by a rapid increase to maximum level at day 10 and sustained thereafter (Fig. 7c). eGFP fluorescence was only seen in cartilaginous nodules starting on day 7 (Fig. 7d). Little eGFP fluorescence was observed in cells that were not chondrogenically induced or ones that had not yet formed cartilaginous nodules. Western blot analysis at day 14 showed that eGFP protein was solely expressed in cells that were exposed to chondrogenic stimuli (Fig. 7e). Taken together, these data illustrate that the 1*cis*CXp promoter has phenotype-specificity similar to the endogenous *Col10a1* promoter, that its transcriptional activity is limited to differentiated ATDC5 cells, and that activity of the gene circuit increases as these cells transition to the hypertrophic phenotype.

**Figure 7.**
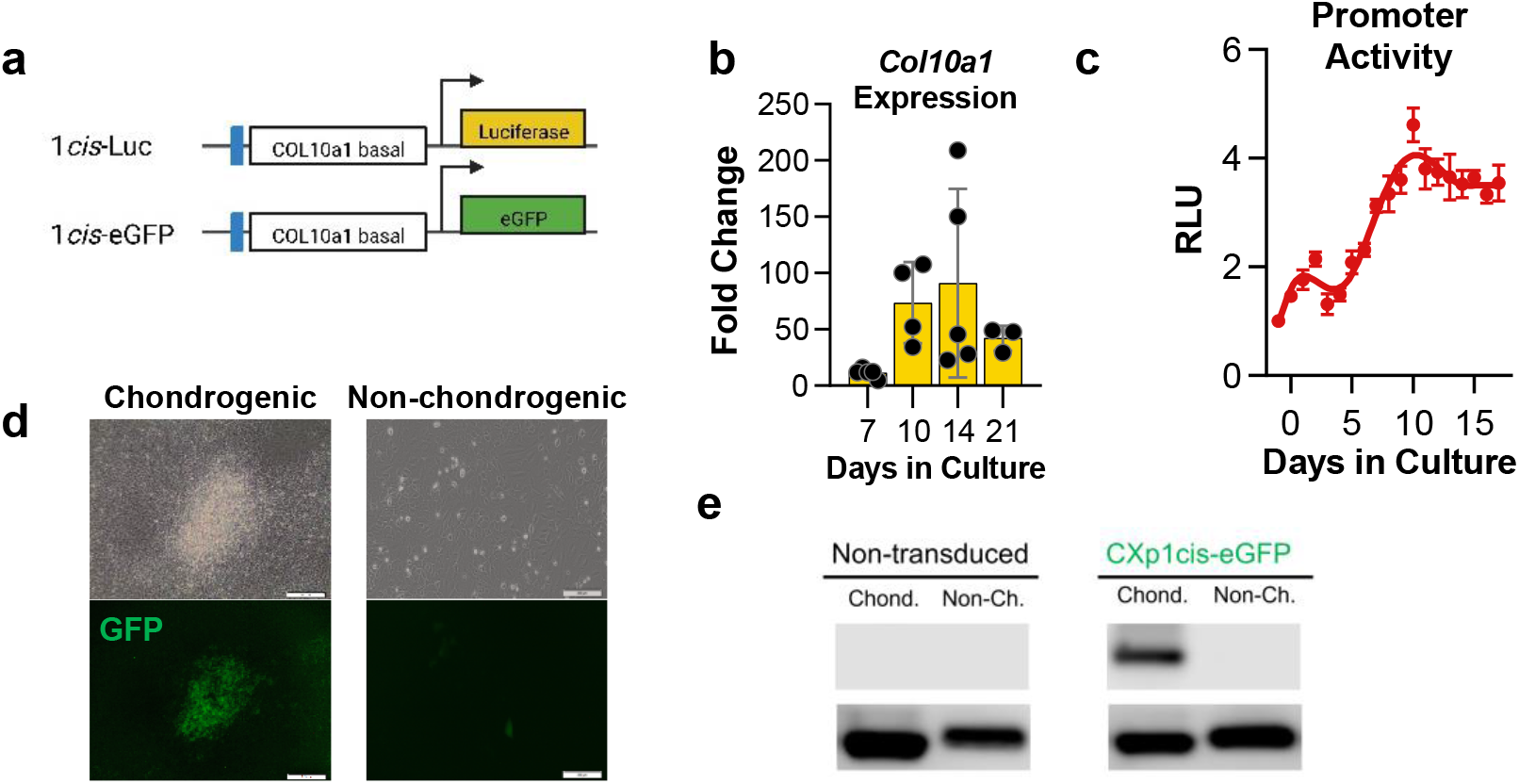
Specificity of the gene circuit. (a) Diagrams of 1*cis*CXp-Luc and 1*cis*CXp-eGFP vectors used to evaluate the activity and chondrogenic specificity, respectively, of the engineered *Col10a1*-like promoter. (b) Quantification of mRNA expression of *Col10a1* in non-transduced ATDC5 monolayer cultures over 21 days of chondrogenic differentiation (two independent experiments, n ≥ 5). (c) Luciferase activity from ATDC5 cultures expressing 1*cis*-Luc over 21 days of differentiation relative to day −1 (three independent experiments, n = 6). (d) Bright field and fluorescent imaging of ATDC5 cells transduced with 1*cis*CXp-eGFP showing that GFP is localized to cartilage nodules of chondrogenic cultures at the onset of *Col10a1* mRNA upregulation (day 7), demonstrating the specificity of the gene circuit activity to chondrogenic cultures. (e) Western blot of eGFP and β-actin (internal control) levels after 14-day chondrogenic cultures in non-transduced ATDC5 cells compared to cells transduced with 1*cis*CXp-eGFP, also demonstrating specificity of the gene circuit activity.

### *Col10a1*-like promoter can be engineered to tune the activity of RUNX2 during chondrogenesis

To determine whether the *Col10a1*-like promoter could be used to provide self-regulated silencing of RUNX2 and to test the hypothesis that the level of RUNX2 suppression could be tuned by varying the number of *cis* enhancers, we monitored the activity of *cis*CXp-sh*Runx2* gene circuits during the chondrogenesis of polyclonal ATDC5 cell populations stably expressing these vectors (Fig. 3). We similarly synthesized 2*cis*-sh*Runx2* and 3*cis*-sh*Runx2* gene circuits and their corresponding scramble control vectors. During the 21-day monolayer chondrogenic differentiation assay, luciferase activity was used as a surrogate measurement of promoter activation.

Luminescence was minimal in cells expressing the 1*cis*-sh*Runx2* vector during the first 6 days, similar to cells transduced with 1*cis*-scramble vector. From day 7, *shRunx2* containing cells exhibited a significantly lower level of luminescence than the scramble controls (Fig. 8a and b). While the total luminescence from both groups fluctuated throughout further differentiation, a stable relative decrease in RUNX2 activity of 18.1 ± 5.1% compared to scrambled controls was reached under the 1*cis*-sh*Runx2* vector. Similarly, gene circuits containing the 2*cis*CXp promoter exhibited the same level of minimal activity as 1*cis*CXp during the first three days of chondrogenic differentiation. Thereafter, the 2*cis*CXp promoter drove the total gene circuit activity in 2*cis*-sh*Runx2* cultures to equilibrate at a level that was 30.4 ± 4.2% lower than the scrambled controls. The addition of a third *cis*-enhancer exhibited a more prominent reduction (77.5 ± 2.6%) in gene circuit activity starting at the onset of chondrogenesis. Gene expression of *Col10a1* was significantly downregulated only in 3*cis*-sh*Runx2* expressing cultures at D14 (Fig. 8c). Significantly more sGAG was accumulated by 2*cis*-sh*Runx2* and 3*cis*-sh*Runx2* cultures at day 14 (Fig. 8d). Alcian blue stained sections showed an increase in the staining area of cartilage nodules. RUNX2 protein expression decreased with increasing numbers of *cis*-enhancers compared to the scramble control (Fig. 8e).

**Figure 8.**
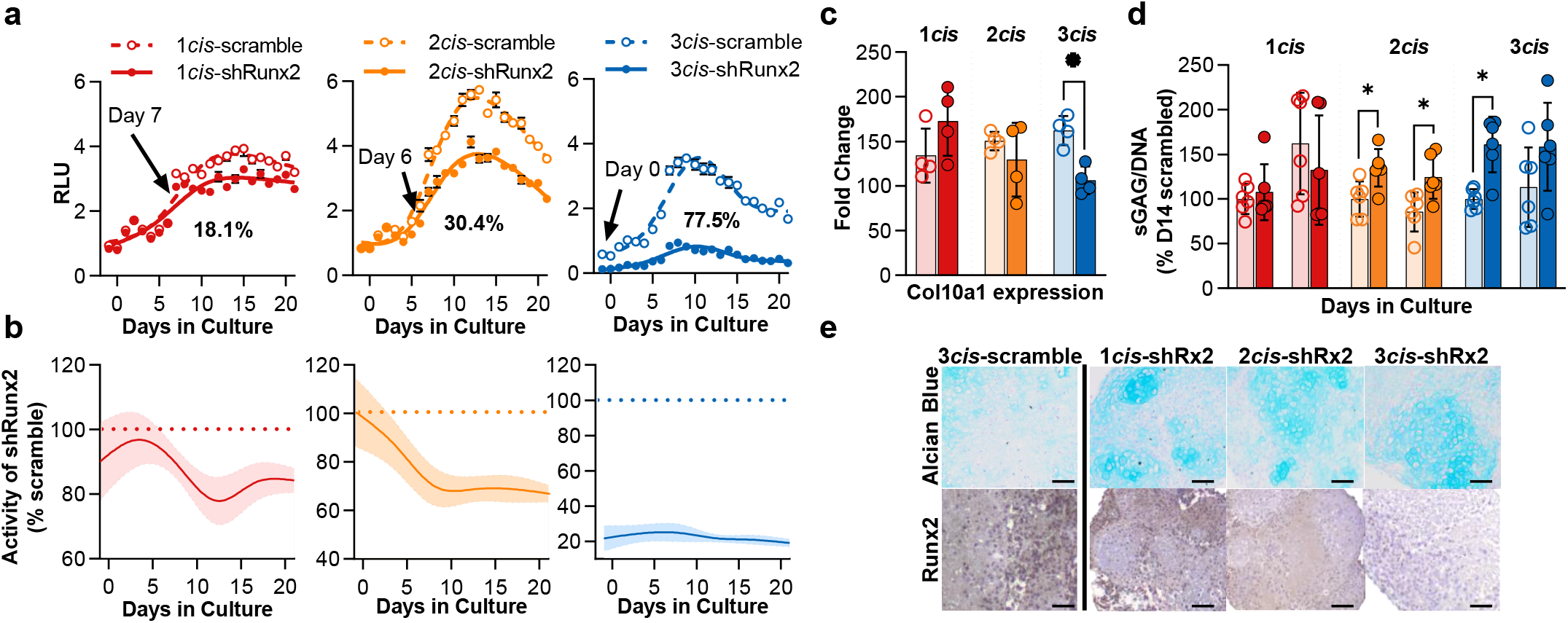
Activity of RUNX2 Suppressing Gene Circuits and Their Effect on Chondrogenesis. (a) Luminescence measured over 21-day chondrogenic cultures expressing 1*cis*CXp-sh*Runx2*/scramble, 2*cis*CXp-sh*Runx2*/scramble, 3*cis*CXp-sh*Runx2*/scramble as a measure of RUNX2 and gene circuit activity. (b) Relative activity of each gene circuit was calculated by normalizing to activity of corresponding scramble controls at each time point. (c) Quantification of mRNA expression (relative to day 0 and the housekeeping genes *Hprt* and *Ppia*) of the RUNX2 target gene, *Col10a1*, at day 14. (d) Fold change in sGAG accumulation normalized to DNA content at day 14 and 28. (e) Staining for sGAG and Runx2 in ATDC5 pellet cultures after 28 days of culture. Data represented as mean ± SD. Significant difference scramble (open circles) and shRunx2 (closed circles) at each time point (n=6 from two independent transduction experiments) is indicated by * p<0.05.

### RUNX2 suppressing gene circuits attenuate hypertrophy in MSC-derived chondrocytes

We next evaluated the function of the gene circuit in human mesenchymal stem cells (hMSCs) during chondrogenesis and neo-cartilage formation. Alcian blue-positive matrix was observed in all pellets created from hMSC-derived chondrocytes (MdChs) expressing *cis*CXp-shRunx2 gene circuits after 28-days in chondrogenic culture. The peripheral regions of the pellets of modified cells had more intense Alcian blue staining, accompanied by less hypertrophy-like cell morphology compared to MdChs derived from WT hMSCs (Fig S3). In response to hypertrophy-inducing stimuli, hMdChs reprogrammed with *cis*CXp-shRunx2 gene circuits accumulated more cartilage matrix compared to both WT and scramble controls (Fig. 9). The *cis*CXp-shRunx2 gene circuit also suppressed mineral deposition under hypertrophic conditions (Fig. 9b). Taken together, this data indicates that RUNX2 suppressing gene circuits are successfully able to prevent hypertrophic maturation of hMdChs and increase their matrix accumulation *in vitro*.

**Figure 9.**
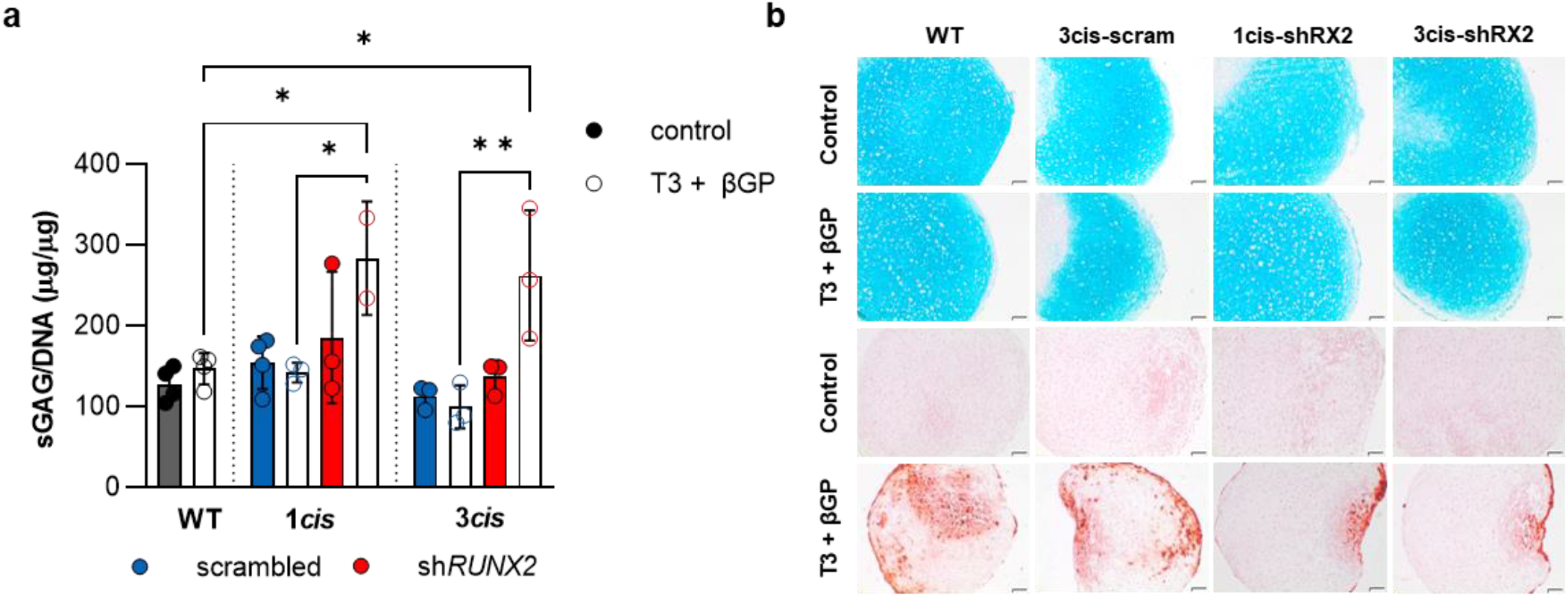
RUNX2 Suppressing Gene Circuits attenuate hypertrophy in human MSC-derived chondrocytes. (a) sGAG accumulation by hMdChs at day 21 after 7 days of hypertrophic induction (b) Alcian blue staining for sGAG and alizarin red staining for calcium deposition in untreated and T3 + βGP treated hMdCh pellets. Scale bar: 50μm.

### RUNX2 suppressing gene circuits mitigates the catabolic effects of IL-1β on MdCh cartilage formation

RUNX2 activity is also upregulated in hMdChs in response to inflammatory stimulus^29–31^. Therefore, we evaluated the effect of RUNX2 suppression on hMdChs response to IL-1β treatment using the 3*cis*-sh*RUNX2* gene circuit to induce maximum RUNX2 suppression in hMdChs. Within 6 hours of the addition of IL-1β to hMdCh cultures, the levels of luminescence, and thus gene circuit activity, were significantly elevated in 3*cis*-scramble pellets and remained unchanged in 3*cis*-sh*RUNX2* pellets (Fig. 10a). There was higher retention of sGAG in 3*cis*-sh*RUNX2* modified hMdCh pellets compared to WT and scrambled controls (Fig. 10b).

**Figure 10.**
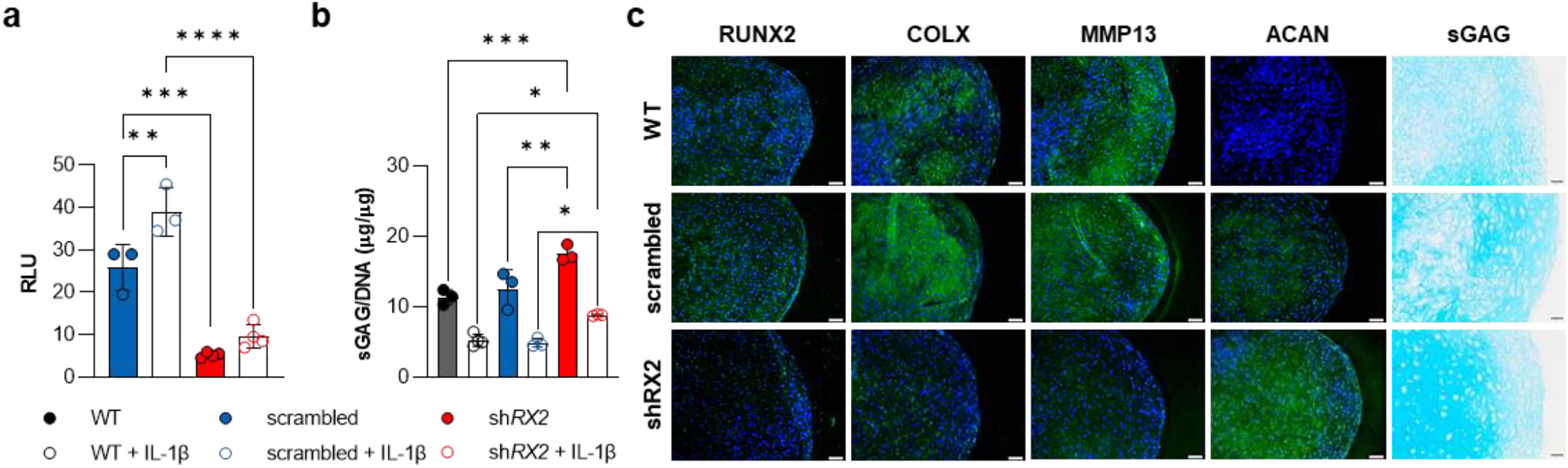
Effect of RUNX2-suppressing gene circuits on MdCh response to IL-1β. (a) Luminescence measured in pellet cultures of hMSC expressing 3*cis*CXp-sh*Runx2*/scramble, after 6 hours of 0.1ng/ml IL-1β treatment (n=3). **(b)** sGAG accumulation normalized to DNA content and **(c)** Alcian blue staining for sGAG and immunofluorescence staining for hypertrophic (RUNX2, COLX, MMP13) and chondrogenic marker (ACAN) in WT and 3*cis*CXp-sh*Runx2*/scramble MdCh pellets treated with 0.1ng/ml IL-1β. Scale bar 50μm

Immunofluorescence analysis revealed lower expression of hypertrophic proteins RUNX2, COLX and MMP13 as well as elevated expression of aggrecan and more intense sGAG staining in 3*cis*-sh*RUNX2* modified hMdCh pellets (Fig. 10c). Taken together, these results show that RUNX2 suppression in hMdCh pellets partially rescues hMdChs from IL-1β induced cartilage matrix catabolism, potentially by reducing the expression of matrix degrading enzymes like MMP13 that are downstream targets of RUNX2 activity.

## DISCUSSION

Here, we demonstrate that the loss of RUNX2 function during chondrogenesis of mesenchymal progenitors can elicit distinct cellular responses at different stages of differentiation. Using a 2D ATDC5 model with inducible *shRunx2* expression, we show that RNAi of *Runx2* in the undifferentiated cells inhibits mesenchymal proliferation and differentiation into chondrocytes. However, induced RNAi of *Runx2* after the pre-chondrogenic proliferation phase or early chondrogenesis enhances the accumulation of cartilaginous matrix structural macromolecules. We further demonstrate that the synthetic *Col10a1*-like promoters can initiate RNAi of *Runx2* in chondrocytes that are transitioning to the pre-hypertrophic phenotype in response to the increasing intracellular RUNX2 activity without interfering with early chondrogenesis. The induced loss of RUNX2 function in turn negatively regulates the activity of the *cis*-sh*Runx2* gene circuit to resist upregulation of RUNX2 during hypertrophy and maturation-associated matrix degradation. Together, our findings highlighted three key features of the *cis*-sh*Runx2* gene circuit: 1) phenotype-specific activation of RNAi, 2) closed-looped, intracellular negative-feedback regulation of RUNX2 activity, and 3) tunable levels of RUNX2 repression. We show that these features of the gene circuits can enhance accumulation of cartilaginous matrix by MdChs during neo-cartilage formation under conditions that drive loss of cartilage tissue functionality in the injured joint, namely hypertrophy and inflammation.

### RUNX2 is essential to chondrogenesis and plays different roles at different stages

Unlike the well-established role of RUNX2 in driving chondrocyte maturation, its involvement in early events of chondrogenesis remains unclear. Progenitor cells within the pre-chondrogenic condensation first proliferate before committing to chondrogenic differentiation^32–34^ and *Runx2* expression has been detected within these condensations^35–37^. Recently, Dexheimer *et al*. showed that this transient phase of proliferation is required for *in vitro* chondrogenesis of human MSCs after cells are condensed into a micromass pellet^38^. In our study, when chondrogenesis is induced in confluent monolayer ATDC5 cultures, differentiation initiates with a similar phase of proliferation that overrides contact inhibition during the first 4 days, which does not occur under the RNAi of *Runx2* (Fig S2). In accordance with our observations, Akiyama *et al*. also showed that introduction of the dominant negative form of RUNX2 in ATDC5 cells inhibits cellular condensation and subsequent chondrogenic differentiation^39^. Similar suppression of chondrogenesis is also observed when *Zfp521*, an inhibitor of RUNX2, is overexpressed in ADTC5 cells^40^. However, it is worth noting that *Runx2^-/-^* mice do form cartilaginous skeleton^41,42^. Nonetheless, prior *in vitro* studies and ours show that constitutively active RNAi of *Runx2* is not suitable cartilage tissue engineering possibly due to the redundant functions of other RUNX proteins and the complexity of the *in vivo* system compared to cells in isolation under static, *in vitro* cultures.

As chondrogenesis proceeds, the role of RUNX2 changes^43^. Delayed RUNX2 silencing in chondroprogenitors after the pre-chondrogenic proliferation phase does not interfere with their further chondrogenic progression; instead it increases the amount of matrix accumulated by these differentiated chondrocytes and decreases the gene expression of MMP13. After pre-chondrogenic proliferation *in vivo*, Nkx3.2-mediated repression of *Runx2* promotes early chondrogenesis by activating *Sox9*^44^. These findings are analogous to our observation of the transient upregulation of RUNX2 prior to the elevated levels of *Col2a1* and *Acan* expression (Fig S1). They also likely explain why loss of RUNX2 function after day 4 no longer blocks chondrogenesis in ATDC5 cells. As in growth plate chondrocytes^45^, the low protein expression of RUNX2 is not permanent in the ATDC5 model. Quickly following the upregulation of *Col2a1* and *Acan*, protein expression of RUNX2 rose and drove differentiated chondrocytes to the pre-hypertrophic and then hypertrophic phenotypes. As expected, induced RUNX2 silencing at this stage of chondrocyte maturation decreased mRNA levels of both *Col10a1* and *Mmp13*. Since both aggrecan and type II collagen are degradation targets of MMP13^46–48^, this outcome is consistent with our observation that RUNX2 silencing leads to an increased level of matrix accumulation under hypertrophic as well as inflammatory environments.

An important observation from the inducible study is that RNAi of *Runx2* in chondrocytes that are transitioning to the pre-hypertrophic phenotype maximizes the accumulation of matrix since it is a net function of production and turnover. Therefore, maximizing the accumulation of cartilaginous matrix via RNAi of *Runx2* requires balancing the inhibition of degradation and increasing ECM production by optimizing the timing and dosing of sh*Runx*2 expression. While RUNX2 silencing is desirable for reducing MMP13-mediated matrix degradation, we also noticed that it also downregulated the gene expression of *Acan* and *Col2a1* to varying levels. While the relationship RUNX2 activity and transcription of *Acan* and *Col2a1* remains to be clarified, RUNX2, together with RUNX1, has been shown to induce the expression of *Sox5* and *Sox6*, which further control the induction of *Col2a1*^49^. Additionally, RUNX2 is a common target of TGF-β1 and BMP-2, both of which are frequently used to induce chondrogenesis of MSCs^50^. These studies, together with our results, suggest that RUNX2 activity may also contribute to the production of aggrecan and type II collagen. Although we cannot easily decouple the regulation of early chondrogenic markers and matrix-degrading enzymes by RUNX2, the differential expression profiles of early and late chondrogenic markers allow us to optimize the temporal activation of sh*Runx2* expression to maximize the accumulation of matrix during chondrogenesis. Identifying such an optimal time is critical because the premature loss of RUNX2 function reduces the production of aggrecan and type II collagen while the belated silencing permits uncontrolled matrix degradation.

### The *cis*CXp-shR*unx2* gene circuit is designed to enable autonomous RUNX2 suppression in heterogeneous stem cell population in preparation for unknown environmental cues

The *cis*CXp-sh*Runx2* gene circuit relies on its synthetic *Col10a1*-like promoter to induce hypertrophy-specific RUNX2 silencing. The specific expression of *Col10a1* in pre-hypertrophic and hypertrophic chondrocytes requires the binding of RUNX2 in addition to the recruitment of the general transcription factors near the transcription start site^26,51,52^, which usually reside within the basal region of mammalian polymerase II promoters. The 330-bp *Col10a1* basal promoter we incorporated contains a highly conserved sequence that precisely describes the transcription start site of *Col10a1* across species^26^, providing the DNA template that supports the assembly of the RNA polymerase II transcription initiation complex. Meanwhile, the two putative tandem-repeat within the 150-bp *cis*-enhancer ensures direct binding of RUNX2^26,53^. As a result of the cooperative actions of these two regulatory elements, all three versions of *cis* promoter are sufficient to direct hypertrophy-specific transcription resembling the endogenous *Col10a1* promoter in differentiating chondrocytes. Critically, the suppression of early chondrogenesis can be avoided as the loss of RUNX2 function does not occur until progenitors fully differentiate into chondrocytes and transition to pre-hypertrophy.

### The *cis*CXp-sh*Runx2* gene circuits establish closed-loop intracellular negative-feedback regulation of RUNX2, allowing differentiating chondrocytes to dynamically resist maturation in response to exogenous cues

The negative feedback motif of *cis*CXp-sh*Runx2* utilizes RUNX2 activity as the central signal. In chondrocytes that are transitioning to hypertrophy, the *cis*CXp promoter initiates the production of *shRunx2* that downregulates RUNX2, which in turn decreases the transcriptional activity of the *cis*CXp promoter. The negative feedback regulation of chondrocyte maturation also occurs naturally, often limited by the range of paracrine signaling (e.g., PTHrP/IHH feedback loop)^54–56^. Free from the dependence on paracrine signaling, *cis*CXp-sh*Runx2* gene circuits allow chondrocytes to resist maturation based their internal tendency to undergo hypertrophy, measured by RUNX2 activity. As a gatekeeper, RUNX2 mediates the signaling of many molecular and biophysical cues in chondrocytes^57,58,59^. Therefore, by inputting the intracellular RUNX2 activity as the signal for the negative-feedback regulation, cells expressing *cis*CXp-sh*Runx2* can dynamically adjust the level of silencing required to maintain their steady states (levels of RUNX2 suppression) without needing to address the specific pathways underlying hypertrophy stimuli, such as thyroid hormone T3 and beta glycerophosphate or inflammatory factors like IL-1β (Fig. 4). Finally, design of *cis*CXp promoter allows *cis*CXp-sh*Runx2* gene circuits to provide a tunable level of RUNX2 suppression. We observed that increasing the number of *cis*-enhancers in the gene circuit progressively enhanced RUNX2 silencing from 18% to nearly 80%. Increasing the number of *cis-*enhancers also led to faster equilibrium of the activity of the gene circuits.

## CONCLUSION

Overall, our circuit design is modular with a straightforward construction, which allows for simple testing of various combinations of circuit components and outputs. This flexible design enables circuits to be optimized for a variety of environmental conditions. Our results demonstrate that this approach could be a viable alternative to strategies that use broad spectrum spatial/temporal regulation of exogenous cues or constitutive or transient silencing of intracellular messengers. However, the behavior of *cis*CXp-sh*Runx2* or similar gene circuits need to be further investigated in long-term *in vitro* and *in vivo* studies. Future studies in the gene circuit design should focus on fine tuning promoter activity to specific cell states, reducing off target effects, and targeting transcriptional and translational regulators of cell function.

## ACKNOWLEDGEMENTS

We thank Ciara Davis, John Braford, Meghan Burns, Hannah Floyd, Samad Emory, and Sunny Karnan for their assistance in analysis of the data and Colleen Flanagan for her editorial assistance. Supported by the ANRF Award No. 310 047420, Orthopaedic Research and Education Foundation Award No. 651100, and NIH-NIAMS Award No. 1R21AR07401101.

## SUPPLMENTAL MATERIALS

**Table S1.**
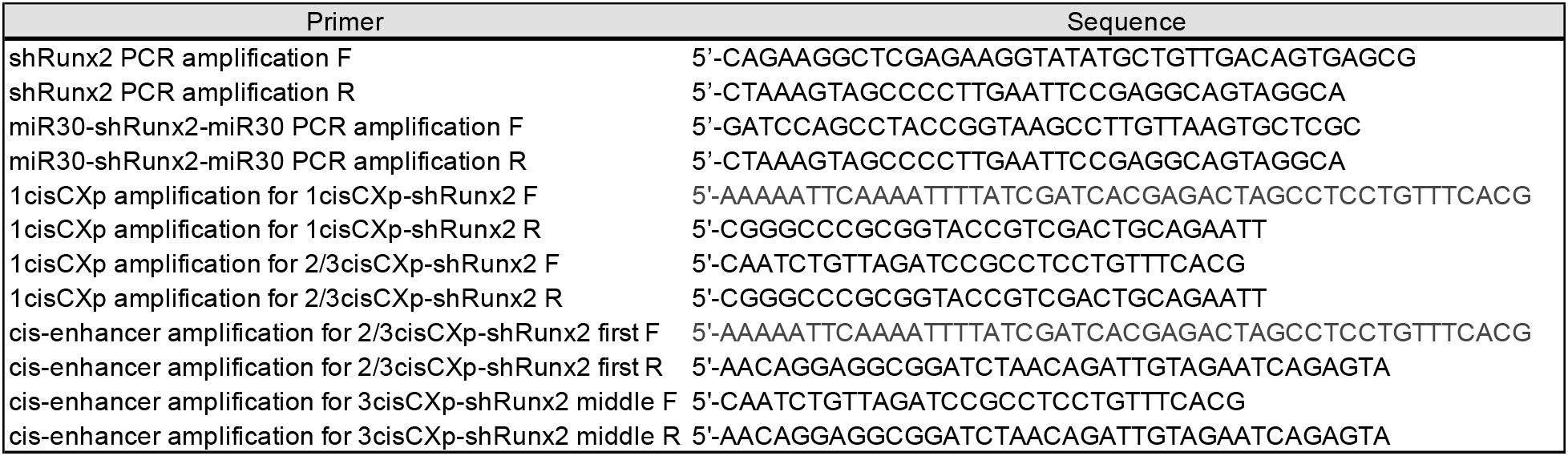
Primers for vector cloning.

**Table S2.**
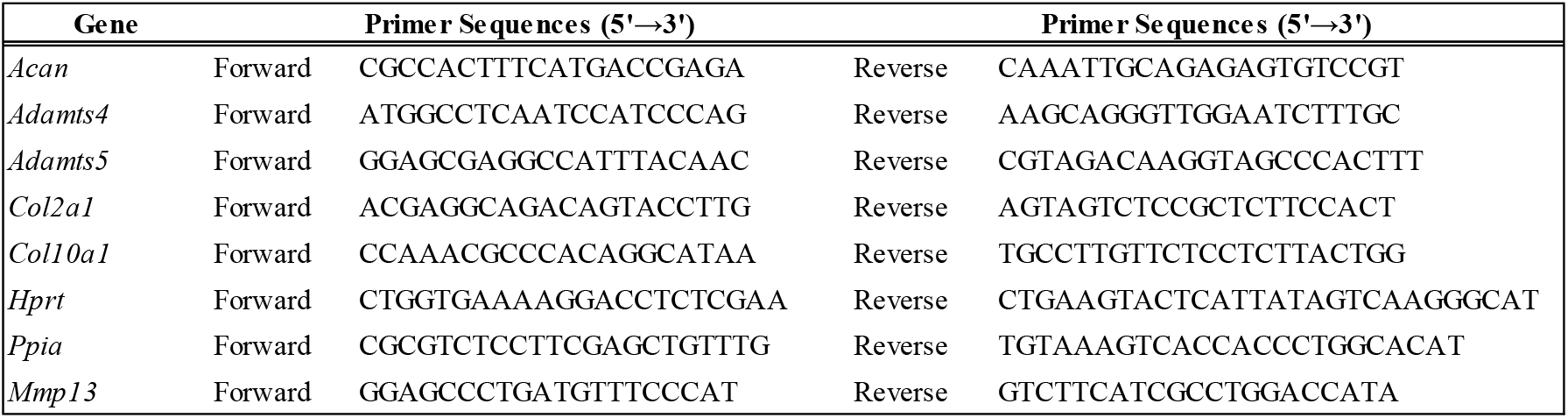
qPCR primers.

**Figure S1.**
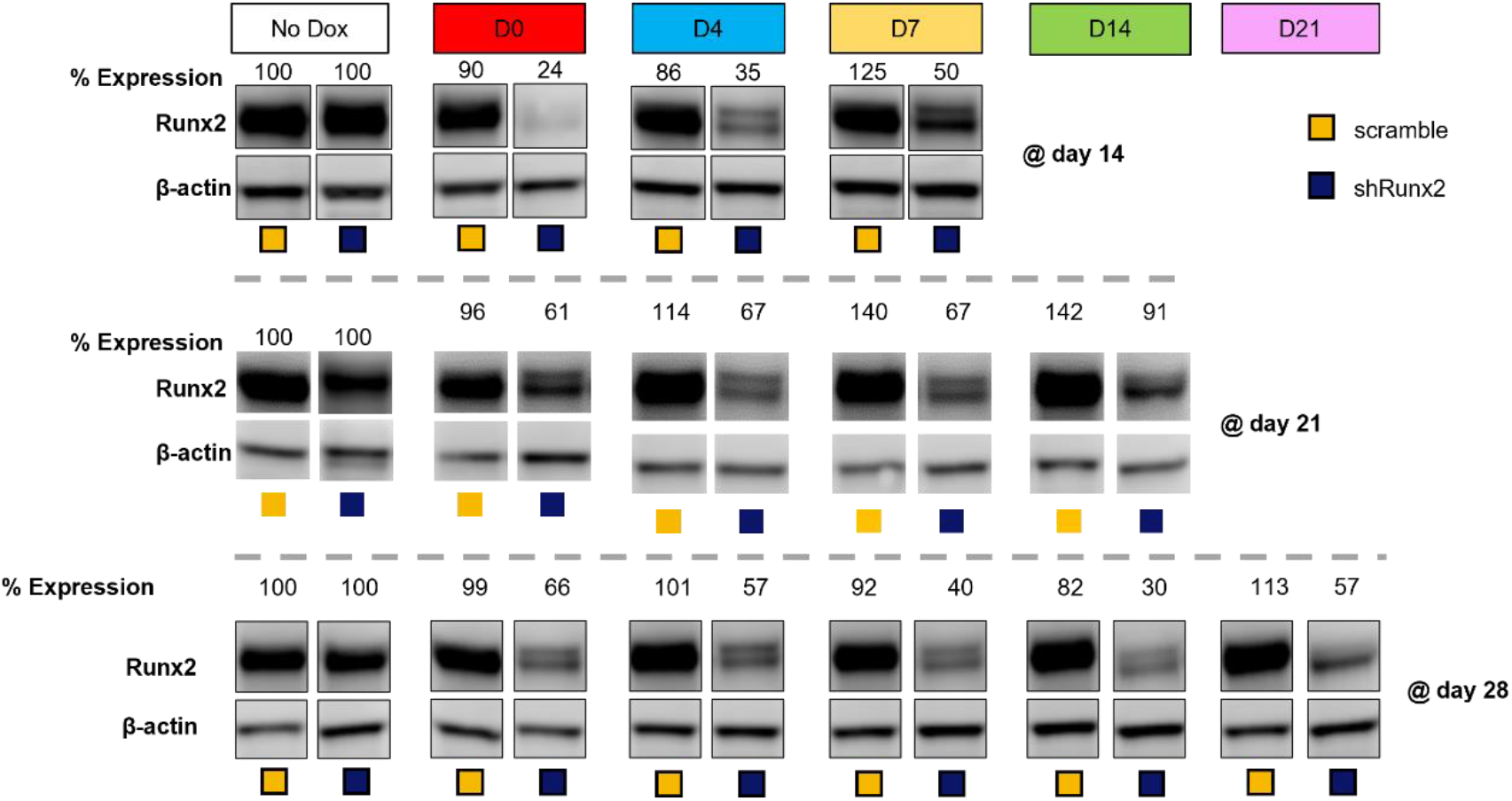
Time-course of Runx2 protein expression. ATDC5 cells containing the inducible gene circuit were treated with doxycycline beginning on the day list across the top (D0, 4, 5, 14, or 21) and continued until Western blot analysis at the time-points indicated (D14, 21, or 28).

**Figure S2.**
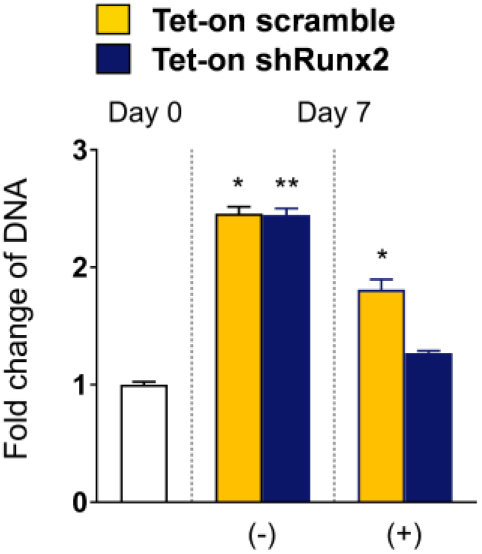
Fold change (relative to day 0) in DNA content at day 7 (different groups at day 7 were compared against day 0)

**Figure S3.**
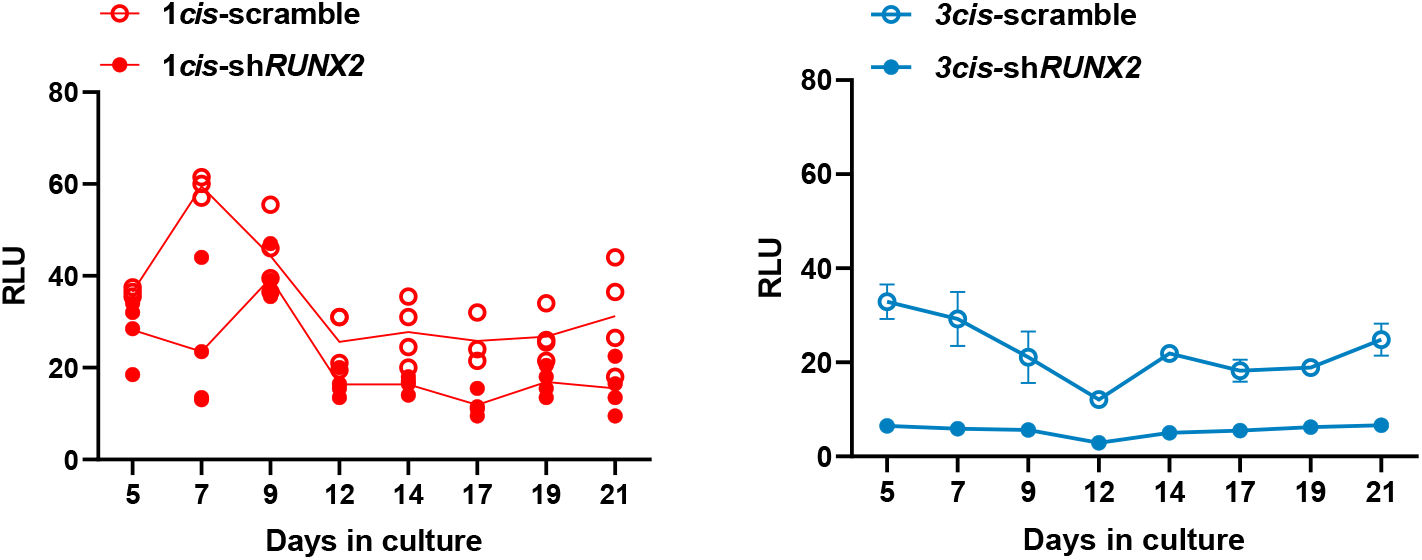
Activity of RUNX2 gene circuits in human MSC derived chondrocytes. Luminescence measured in 21-day chondrogenic cultures expressing 1*cis*CXp-sh*Rwnx2*/scramble, 3*cis*CXp-sh*Runx2*/scramble as a measure of RUNX2 activity.

**Figure S4.**
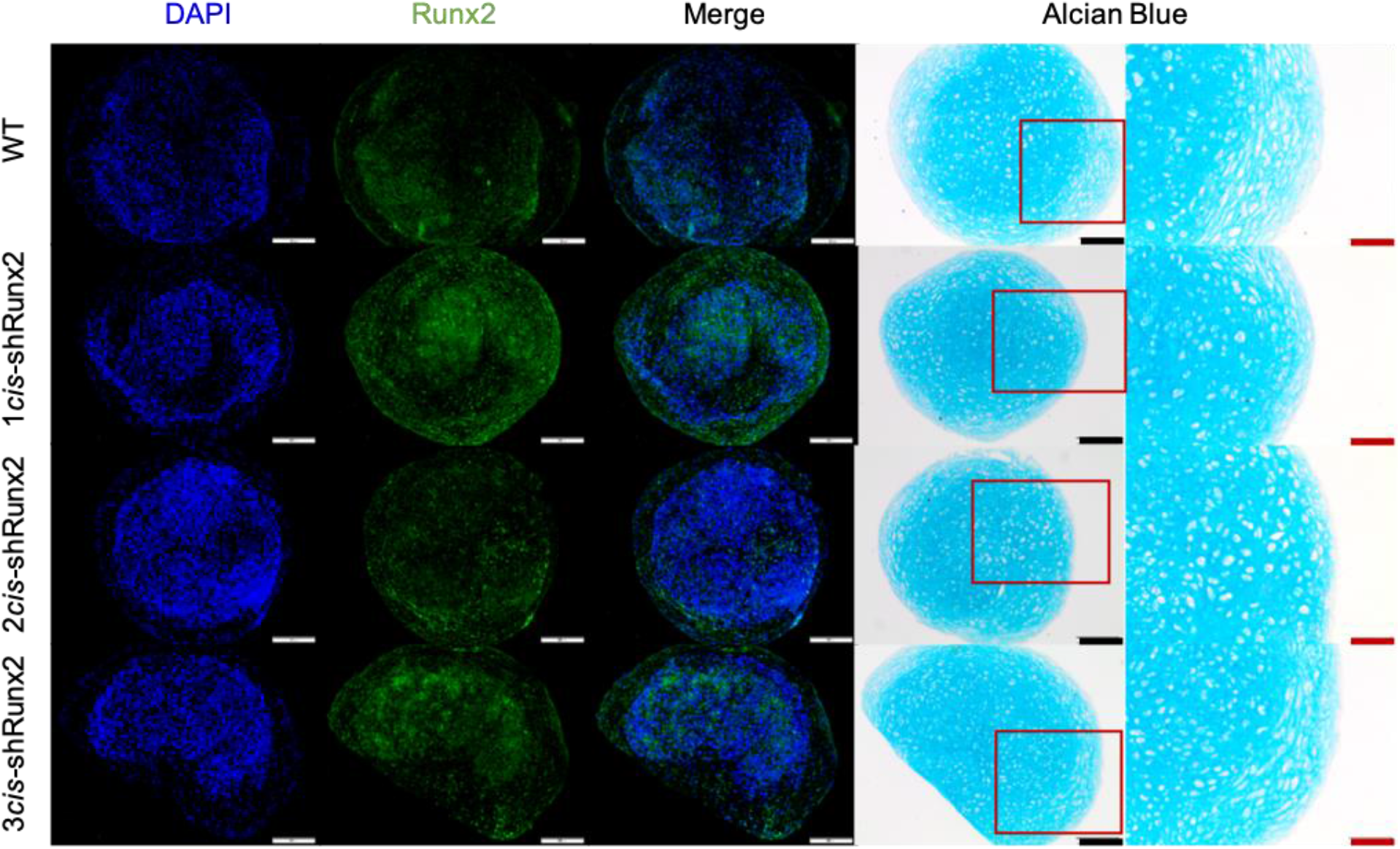
Immunoflourescent staining for RUNX2 protein expression and the corresponding Alcian Blue staining of chondrogenic pellets derived from WT and *cis*CXp-sh*Runx2* hMSC at day 28 of chondrogenic culture.

