## Supplement materials for "A Synthetic, Closed-Looped Gene Circuit for the Autonomous Regulation of RUNX2 Activity during Chondrogenesis"

### SUPPLMENTAL MATERIALS

#### Table S1. Primers for vector cloning

| Primer | Sequence |
|---|---|
| shRunx2 PCR amplification F | 5’-CAGAAGGCTCGAGAAGGTATATGCTGTTGACAGTGAGCG |
| shRunx2 PCR amplification R | 5’-CTAAAGTAGCCCCTTGAATTCCGAGGCAGTAGGCA |
| miR30-shRunx2-miR30 PCR amplification F | 5’-GATCCAGCCTACCGGTAAGCCTTGTTAAGTGCTCGC |
| miR30-shRunx2-miR30 PCR amplification R | 5’-CTAAAGTAGCCCCTTGAATTCCGAGGCAGTAGGCA |
| 1cisCXp amplification for 1cisCXp-shRunx2 F | 5'-AAAAATTCAAAATTTTATCGATCACGAGACTAGCCTCCTGTTTCACG |
| 1cisCXp amplification for 1cisCXp-shRunx2 R | 5'-CGGGCCCGCGGTACCGTCGACTGCAGAATT |
| 1cisCXp amplification for 2/3cisCXp-shRunx2 F | 5'-CAATCTGTTAGATCCGCCTCCTGTTTCACG |
| 1cisCXp amplification for 2/3cisCXp-shRunx2 R | 5'-CGGGCCCGCGGTACCGTCGACTGCAGAATT |
| cis-enhancer amplification for 2/3cisCXp-shRunx2 first F | 5'-AAAAATTCAAAATTTTATCGATCACGAGACTAGCCTCCTGTTTCACG |
| cis-enhancer amplification for 2/3cisCXp-shRunx2 first R | 5'-AACAGGAGGCGGATCTAACAGATTGTAGAATCAGAGTA |
| cis-enhancer amplification for 3cisCXp-shRunx2 middle F | 5'-CAATCTGTTAGATCCGCCTCCTGTTTCACG |
| cis-enhancer amplification for 3cisCXp-shRunx2 middle R | 5'-AACAGGAGGCGGATCTAACAGATTGTAGAATCAGAGTA |

#### Table S2. qPCR primers

| Gene | | Primer Sequences (5'→3') | | Primer Sequences (5'→3') |
|---|---|---|---|---|
| *Acan* | Forward | CGCCACTTTCATGACCGAGA | Reverse | CAAATTGCAGAGAGTGTCCGT |
| *Adamts4* | Forward | ATGGCCTCAATCCATCCCAG | Reverse | AAGCAGGGTTGGAATCTTTGC |
| *Adamts5* | Forward | GGAGCGAGGCCATTTACAAC | Reverse | CGTAGACAAGGTAGCCCACTTT |
| *Col2a1* | Forward | ACGAGGCAGACAGTACCTTG | Reverse | AGTAGTCTCCGCTCTTCCACT |
| *Col10a1* | Forward | CCAAACGCCCACAGGCATAA | Reverse | TGCCTTGTTCTCCTCTTACTGG |
| *Hprt* | Forward | CTGGTGAAAAGGACCTCTCGAA | Reverse | CTGAAGTACTCATTATAGTCAAGGGCAT |
| *Ppia* | Forward | CGCGTCTCCTTCGAGCTGTTTG | Reverse | TGTAAAGTCACCACCCTGGCACAT |
| *Mmp13* | Forward | GGAGCCCTGATGTTTCCCAT | Reverse | GTCTTCATCGCCTGGACCATA |

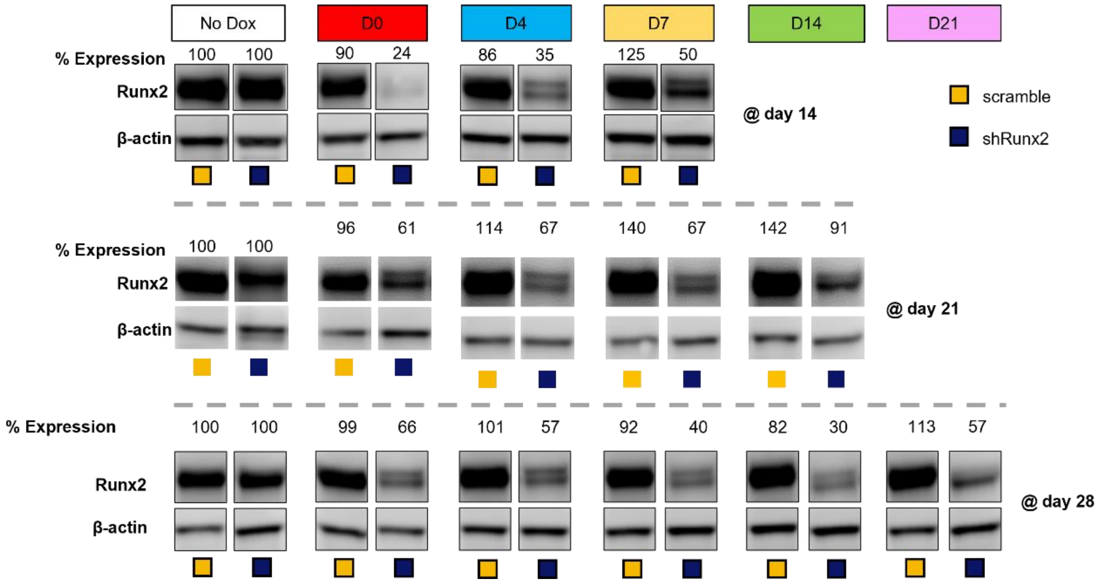


**Figure S1. Time-course of Runx2 protein expression**. ATDC5 cells containing the inducible gene circuit were treated with doxycycline beginning on the day list across the top (D0, 4, 5, 14, or 21) and continued until Western blot analysis at the time-points indicated (D14, 21, or 28).

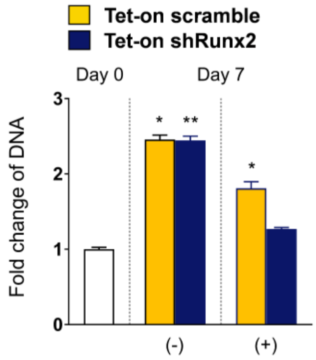


**Figure S2.** Fold change (relative to day 0) in DNA content at day 7 (different groups at day 7 were compared against day 0)

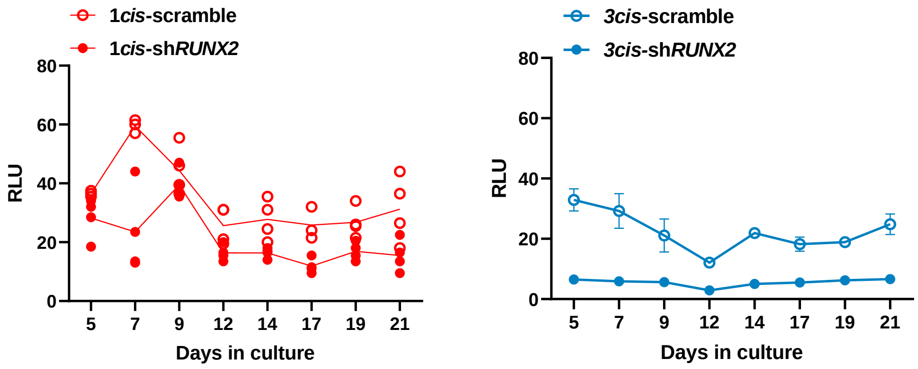


**Figure S3. Activity of RUNX2 gene circuits in human MSC derived chondrocytes.** Luminescence measured in 21-day chondrogenic cultures expressing 1*cis*CXp-sh*Runx2*/scramble, 3*cis*CXp-sh*Runx2*/scramble as a measure of RUNX2 activity.

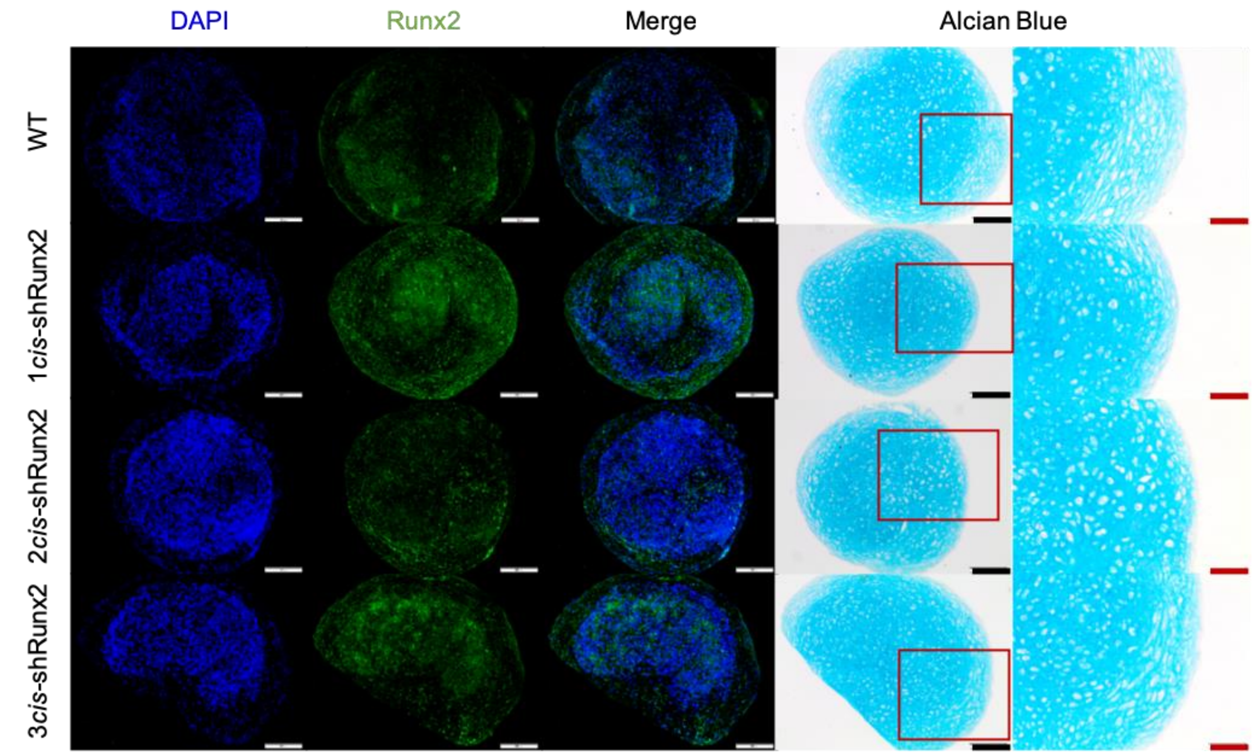


**Figure S4.** Immunoflourescent staining for RUNX2 protein expression and the corresponding Alcian Blue staining of chondrogenic pellets derived from WT and *cis*CXp-sh*Runx2* hMSC at day 28 of chondrogenic culture.
